# Crystal structure and solution state of the C-terminal head region of the narmovirus receptor binding protein

**DOI:** 10.1101/2022.12.02.518945

**Authors:** Alice J. Stelfox, Kasopefoluwa Y. Oguntuyo, Ilona Rissanen, Karl Harlos, Robert Rambo, Benhur Lee, Thomas A. Bowden

## Abstract

Increased viral surveillance has led to the isolation and identification of numerous uncharacterized paramyxoviruses, rapidly expanding our understanding of paramyxoviral diversity beyond the bounds of known genera. Despite this diversity, a key feature that unites paramyxoviruses is the presence of a receptor-binding protein, RBP, which facilitates host-cell attachment and plays a fundamental role in determining host-range. Here, we study the RBP presented on the surface of rodent-borne paramyxoviruses Mossman and Nariva (MosV and NarV, respectively), viruses that constitute founding members of the recently defined *Narmovirus* genus within the *Paramyxoviridae* family. Crystallographic analysis of the C-terminal head region of the dimeric MosV and NarV RBPs demonstrates that while these glycoproteins retain the canonical six-bladed β-propeller fold found in other paramyxoviral RBPs, they lack the structural motifs associated with established paramyxovirus host-cell receptor entry pathways. Consistent with MosV-RBP and NarV-RBP undergoing a distinct entry pathway from other characterized paramyxoviruses, structure-based phylogenetic analysis demonstrates that these six-bladed β-propeller head domains form a singular structural class that is distinct from other paramyxoviral RBPs. Additionally, using an integrated crystallographic and small angle X-ray scattering analysis, we confirm that MosV-RBP and NarV-RBP form homodimeric arrangements that are distinct from those adopted by other paramyxovirus RBPs. Altogether, this investigation provides a molecular-level blueprint of the narmovirus RBP that broadens our understanding of the structural space and functional diversity available to paramyxovirus RBPs.

**Importance:** Genetically diverse paramyxoviruses are united in their presentation of a receptor-binding protein (RBP), which works in concert with the fusion protein to facilitate host-cell entry. The C-terminal head region of the paramyxoviral RBP, a primary determinant of host-cell tropism and inter-species transmission potential, forms structurally distinct classes dependent upon protein and glycan receptor specificity. Here, we reveal the architecture of the C-terminal head region of the RBPs from Nariva virus (NarV) and Mossman virus (MosV), two archetypal rodent-borne paramyxoviruses within the recently established genus *Narmovirus*, family *Paramyxoviridae*. Our analysis reveals that while narmoviruses retain the general architectural features associated with paramyxoviral RBPs, namely a six-bladed β-propeller fold, they lack the structural motifs associated with known receptor-mediated host-cell entry pathways. This investigation indicates that the RBPs of narmoviruses exhibit pathobiological features that are distinct from those of other paramyxoviruses.

## Introduction

The extensive genetic diversity revealed by virus discovery programmes has permitted the creation of new genera within the family *Paramyxoviridae*, including *Narmovirus, Jeilongvirus*, and *Salemvirus* [1-5]. This update to paramyxovirus taxonomy has allowed two previously termed ‘orphan paramyxoviruses’ [6, 7], Nariva virus (NarV) and Mossman virus (MosV), to become founding members of the *Narmovirus* genus. NarV was isolated on four separate occasions in the early 1960s from forest rodents in Trinidad and Tobago [8, 9] and MosV was isolated in the 1970s from the pooled organs of wild rats originating from two separate locations in Queensland, Australia [10, 11]. The ‘bank vole virus’ (BaVV) [12], which was discovered in Russia, and a narmovirus (UKMa K4D) prevalent in approximately four percent of field voles studied in Cheshire and Leicestershire, UK, have been more recently identified and putatively added to this newly established genus [13]. Little is known about the disease burden imposed by these pathogens upon wild animal reservoirs, nor the capacity of these viruses to be transmitted to non-native host species, a characteristic common amongst many paramyxoviruses [14].

An fundamental component of the paramyxovirus life cycle, in both infection of native host reservoirs and during spillover into new host-species, is the ability of the virus to productively interact with host-cell surface receptors during host-cell entry [7]. This process is facilitated by a virus envelope-displayed receptor binding protein (RBP), which is presented on the paramyxovirus surface as a dimer-of-dimers [15]. Each protomer of the RBP consists of an N-terminal intraviral (IV) region, transmembrane (TM) domain, an α-helical stalk, and a C-terminal six-bladed β-propeller head domain, which mediates the interaction with the cognate host-cell receptor [15-18]. The initial interaction between an RBP and receptor precedes virus internalization and fusion glycoprotein (F)-mediated merger of the virus and host-cell membranes [19, 20].

Despite conservation of the overall six-bladed β-propeller fold, paramyxoviral RBPs bind a diverse range of host cell receptors, including carbohydrates and proteins [18, 21, 22]. Viruses within genera *Morbillivirus* and *Henipavirus* are known to use proteinaecous receptors during host-cell entry [7]. Morbilliviruses (e.g., measles virus, MV) present a hemagluttinin (H) RBP, which recognizes both lymphocyte activation molecule-F1 (SLAMF1) and nectin-4 at an overlapping hydrophobic concave groove located at the side of the β4–β6 propeller face [23-27]. Most henipavirus glycoproteins (HNV-G) utilize B-class ephrins as high affinity receptors by interacting at a shallow, yet extensive cleft at the top of the six-bladed β-propeller of the RBP [28-31]. In contrast, paramyxoviruses within the genera *Respirovirus, Orthoavulavirus, Metaavulavirus, Paraavulavirus*, and *Orthorubulavirus*, display RBPs with hemagglutinin-neuraminidase (HN) functionality, which render them capable of recognizing and hydrolyzing sialic acid at the top face of the β-propeller [32-36]. A conserved feature of RBPs with HN functionality is the presence of seven conserved sialidase residues and a conserved hexapeptide motif [33, 36, 37].

Reflective of paramyxovirus RBPs displaying diverse receptor-binding functionality, the primary amino acid sequences of these proteins are highly variable [7]. For example, pair-wise comparison reveals that although MosV-RBP and NarV-RBP cluster together within a phylogenetic tree of RBP sequences [7], their primary sequences exhibit only ∼30% identity. Here, we sought to clarify the structural relationship of MosV-RBP and NarV-RBP glycoproteins with characterized paramyxoviral RBPs. X-ray crystallographic analysis of these RBPs to 1.6 Å and 2.1 Å resolution, respectively, reveals a distinctive six-bladed β-propeller architecture that lacks receptor recognition features associated with H, G, and HN RBPs. Our observed dissimilarities support a model whereby narmoviruses likely use a receptor distinct from known canonical paramyxovirus receptors.

## Results

### Structure determination of functionally active NarV-RBP_β_ and MosV-RBP_β_ RBPs

MosV and NarV have been assigned to the genus, *Narmovirus*, within the family *Paramyxoviridae* [2]. Given the importance of the paramyxoviral RBP in determining cellular and species tropism [7, 21], we sought to ascertain whether the independent classification of NarV and MosV from other paramyxoviruses was reflected in RBP structure. To this end, we produced two sets of soluble constructs of NarV-RBP and MosV-RBP for both structural and functional studies. For structural studies, minimal constructs bearing the C-terminal six-bladed β-propeller head domain of NarV-RBP (Glu184–Pro657, termed ‘NarV-RBP_β_’) and MosV-RBP (Glu202–Thr632, termed ‘MosV-RBP_β_’) tagged with a C-terminal hexahistidine tag were generated (**Fig. 1A**. For cell-binding analysis (**Fig. 1B**), N-terminal Fc tagged soluble constructs of NarV-RBP (Glu184–Pro657, termed ‘Fc-NarV-RBP_β_’) and MosV-RBP (Thr157–Thr632, termed ‘Fc-MosV-RBP_β_’) were produced. Beyond the β-propeller head domain, the NarV-RBP constructs also included a small C-terminal extension of 31 residues of unknown function, which is not present in MosV-RBP and BAVV-RBP and lacks sequence conservation with other paramyxoviral RBPs.

**Figure 1.**
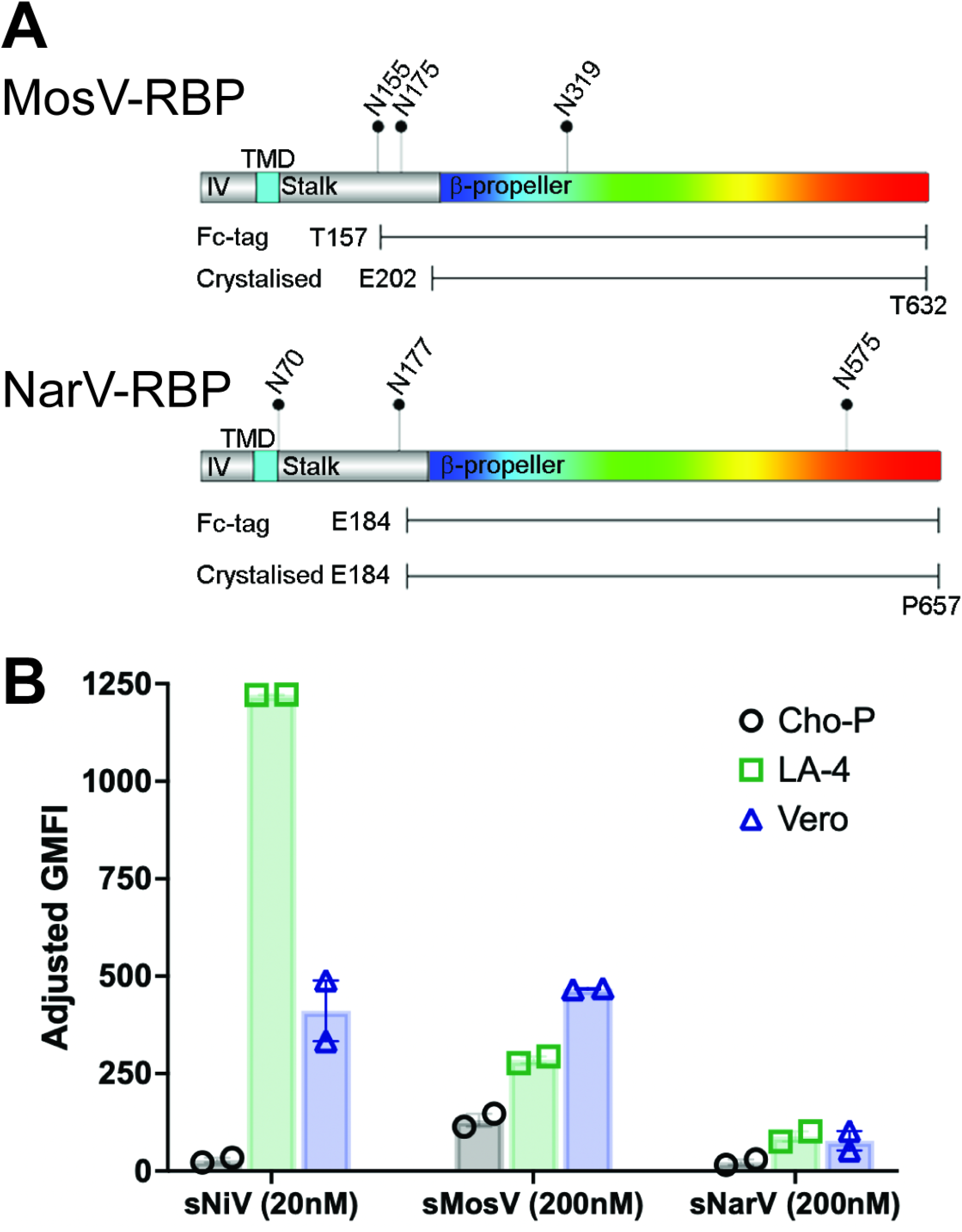
Binding analysis of soluble MosV-RBP and NarV-RBP to a panel of cell-types confirms constructs are functional. (**A**) Gene diagram generated with DOG2.0 displaying features of the MosV and NarV RBPs, including the intraviral (IV, gray) region, transmembrane domain (TMD, cyan), stalk region (gray), and six-bladed β-propeller receptor binding head (rainbow). Predicted N-linked glycosylation sites (NXS/T, where X≠P) are marked with pins and relevant asparagine residue numbered. (**B**) Soluble receptor binding protein (RBP) binding to LA-4, Vero, and CHO pgsA745 cell lines. The indicated concentration of soluble, Fc-tagged RBP was incubated with cells, then stained with APC-tagged anti-Fc secondary antibody and subjected to flow cytometry as described in the methods. Adjusted geometric mean fluorescence intensity (GMFI) was calculated as a product of the percent APC positive cells and the GMFI of the APC positive cells. Shown are the results of two replicates for each cell line with error bars representing the standard error of the mean. MosV-RBP_β_ showed moderate binding and NarV-RBP_β_ showed low level binding to both LA-4 and Vero cells. The decreased binding affinity of NarV-RBP_β_ could be attributed to the shorter construct length.

To evaluate the functionality of NarV-RBP_β_ and MosV-RBP_β_, we measured binding of the purified proteins to Vero (African green monkey) and LA-4 (mouse lung adenoma) cells by flow cytometry. MosV-RBP_β_ and NarV-RBP_β_ bound to both cell types, with MosV-RBP_β_ exhibiting a greater level of overall binding (**Fig. 1B**). These data suggest that both Vero and LA-4 cells display cell-surface receptor(s) recognized by NarV and MosV.

Both NarV-RBP_β_ (Asn575) and MosV-RBP_β_ (Asn319) constructs present a single N-linked glycosylation sequon (as defined by NXT/S, where X≠P) site (**Fig. 1A**). To facilitate crystallogenesis, NarV-RBP_β_ and MosV-RBP_β_ were produced in the presence of the α-mannosidases I inhibitor, kifunensine [38], and the resultant high mannose-type glycans were partially cleaved by endoglycosidase F1 [39]. Crystals of NarV-RBP_β_ and MosV-RBP_β_ diffracted to 2.2 and 1.6 Å resolution, respectively. Neither narmoviral RBP structure was amenable to solution by the molecular replacement method using previously reported paramyxoviral RBP models, which likely reflects the distant structural relationship of NarV-RBP_β_ and MosV-RBP_β_ from other paramyxoviral RBPs. As a result, the MosV-RBP_β_ structure was solved utilizing the single isomorphous replacement with anomalous scattering (SIRAS) method [40] with a platinum derivative, K_2_PtCl_6_ (**Table S1**). Subsequently, the NarV-RBP_β_ structure was phased using the partially built MosV-RBP_β_ structure as a molecular replacement model (**Table S2**).

### The narmoviral RBPs present distinct paramyxoviral receptor binding architectures

Both NarV-RBP_β_ (Thr209–Thr632) and MosV-RBP_β_ (Arg203–Asn626) exhibit the expected six-bladed β-propeller fold common to paramyxoviral RBPs, with each blade composed of four anti-parallel β-strands (**Fig 2A**). The fold is stabilized by seven conserved disulphide bonds, which are also presented by HNV-G RBPs, including NiV-G RBP. With the exception of the terminal residues Glu202 and Thr632, the majority of the MosV-RBP_β_ structure was well-ordered within the crystal. In contrast, more regions could not be resolved in the electron density map of the NarV-RBP_β_ structure, including residues at the N- (Glu184–Asn197) and extended C-termini (Asn626–Pro657), which are directed towards the solvent channels in the crystal. We note that the extended C-terminal regions of Mòjiāng virus G RBP (MojV-RBP) and Ghana virus G RBP (GhV-RBP) were similarly disordered in previous structural studies [29, 41]. These combined observations are suggestive that these unique C-terminal regions of the RBP may be inherently plastic or may require the stabilizing environment of neighboring RBPs or F glycoproteins, as present in the context of the native virion.

**Figure 2.**
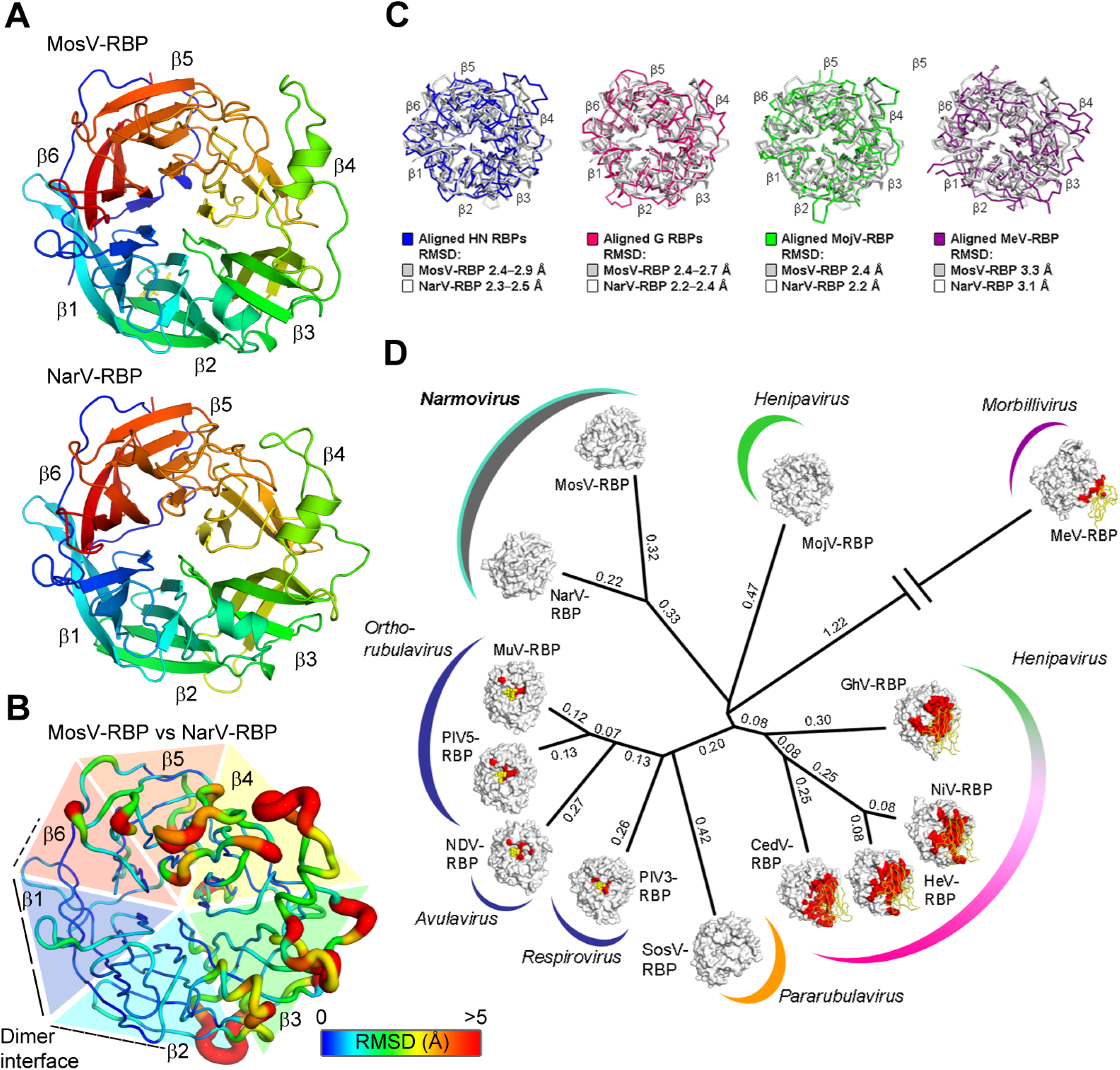
The MosV and NarV RBPs form a singular structural class within the paramyxovirus family. (**A**) The structures of MosV-RBP_β_ and NarV-RBP_β_ are shown in cartoon representation, and colored rainbow from the N- to C-termini (blue to red). The blades of the propeller are labelled β1–β6. (**B**) Calculated root-mean-square deviation (RMSD) between the aligned Cα residues of MosV-RBP_β_ and NarV-RBP_β_ represented through colouration according to RMSD value (blue to red with increasing RMSD), and width of the cartoon (thin to thick with increasing RMSD). RMSD was calculated using secondary structure matching (SSM) [71]. Cα residues that failed to align or had RMSD values of >5 Å, were assigned the value 5 Å. (**C**) Overlays of MosV-RBP_β_ (white) and NarV-RBP_β_ (grey) with other paramyxoviral RBP structures: from left to right, NDV (blue, 1E8V), NiV (pink, 2VWD), MeV (2ZB5), MojV (green, 5NOP). Cα trace rendered and RMSD annotated. (**D**) Clockwise from NarV; MosV; MojV (5NOP); MeV (2ZB5); GhV (4UF7); NiV (2VWD); HeV (2X9M); CedV, Cedar virus (6THB); SosV, Sosuga virus (6SG8); PIV3, parainfluenzavirus 3 (1V2I); NDV (1E8V); PIV5, parainfluenza virus 5; MuV (5B2C). Structural Homology Program (SHP [68]), was used to calculate evolutionary distance matrices by means of pairwise superposition of RBP structures. The resultant matrices were used to plot an unrooted tree in PHYLIP [69]. RBP surfaces are represented with cognate receptor binding sites colored (red), and receptors presented either as yellow ribbon (protein) or spheres (carbohydrate). Calculated evolutionary distances are indicated beside the branches.

Despite exhibiting only 31.4% amino acid sequence identity, NarV-RBP_β_ and MosV-RBP_β_ present similar overall structures, where structural overlay results in a root-mean-square-deviation (RMSD) of 1.6 Å over 401 aligned Cα atoms. Comparative RMSD analysis of the NarV-RBP_β_ and MosV-RBP_β_ (**Fig. 2B**) reveals that higher levels of structural deviation are observed predominantly in blades 4–6 of the β-propeller at solvent exposed loops.

Consistent with the observed genetic distance of narmovirus RBPs from other paramyxovirus RBPs [7], overlay analysis reveals that MosV-RBP_β_ and NarV-RBP_β_ are structurally distinct from measles virus H (MeV-H) RBP_β_ (3.3 Å and 3.1 Å RMSD, respectively), HNV-G RBP_β_ structures (2.4–2.7 Å and 2.2–2.4 Å RMSD, respectively) [42], and HN-type RBP_β_s (2.4–2.9 Å and 2.3–2.5 Å RMSD, respectively) [3, 23, 29, 32-35] (**Fig. 2C**). This observed independence is reflected upon structure-based phylogenetic analysis, which reveals that MosV-RBP_β_ and NarV-RBP_β_ form a distinct branch that is nearly equidistant from the H, G, and HN-type RBPs (**Fig. 2D**). This analysis demonstrates that narmovirus RBPs fall outside established receptor-specific structural groupings.

### Narmoviral RBPs are dimeric in solution and in the crystal

Size exclusion analysis of purified NarV-RBP_β_ and MosV-RBP_β_ revealed that the two proteins form putative dimers in solution (**Supplementary Fig. S1**). Furthermore, the asymmetric unit of both NarV-RBP_β_ and MosV-RBP_β_ structures consists of two near-identical β-propeller head domains, where each protomer forms an extensive protein–protein interface between the first (β1) and sixth (β6) blades of the β-propeller (**Fig. 3**). As calculated by the ‘Proteins Interfaces Surfaces Assemblies’ (PISA) server [43], the association between MosV-RBP protomers occludes ∼2,070 Å^2^ of solvent accessible surface area, and is stabilized by 16 hydrogen bonds and 3 salt bridges. The interface between NarV-RBP protomers is similarly substantial, with an occluded surface area of ∼2,160 Å^2^ stabilized by 19 hydrogen bonds and 3 salt bridges.

**Figure 3.**
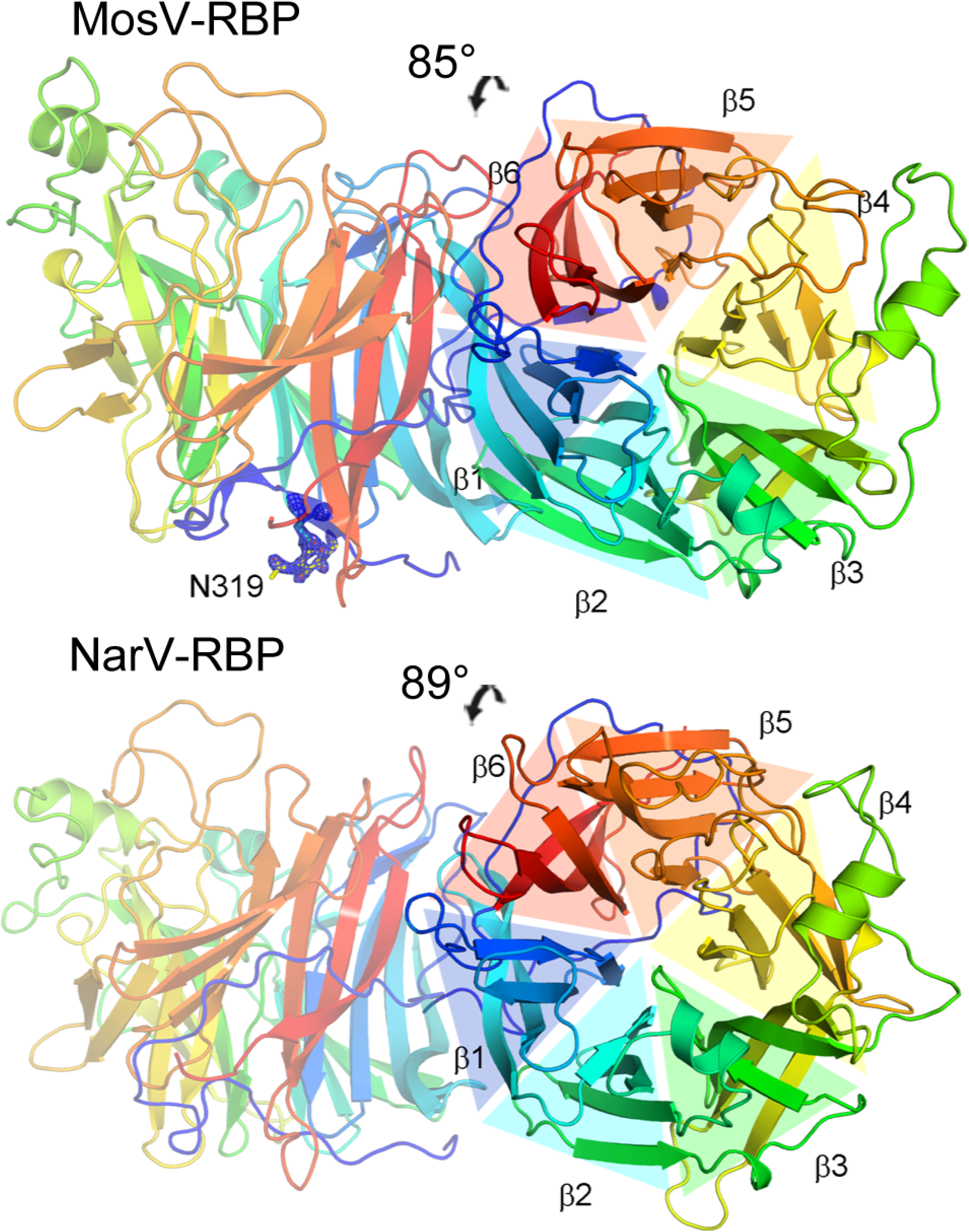
MosV-RBP_β_ and NarV-RBP_β_ present homodimeric interfaces. The crystallographic MosV-RBP and NarV-RBP dimer shown in cartoon representation with monomers colored rainbow from the N-terminus to C-terminus (blue to red, respectively). The calculated angle of association is shown above the dimer.

Interestingly, the mode of MosV-RBP and NarV-RBP dimerization contrasts that observed in previously reported H, HN, and G-type RBP structures. Indeed, although all reported homodimeric RBP structures utilize the first and/or sixth blades (β1 and β6, respectively) of the β-propeller, the narmovirus RBP utilizes a different angle of association and level of buried surface to previously reported paramyxovirus RBP dimers. Indeed, while the HN-type RBP dimers typically assemble with a 60° angle of association between protomers, with ∼1,790 Å^2^ of buried surface [37], the MosV-RBP and NarV-RBP protomers are organized approximately ∼90° relative to each other (**Fig. 3**). This mode of dimerization also contrasts morbillivirus MV-H (∼120° and ∼1,080 Å^2^, respectively) and henipavirus HeV-G RBP (∼40° and ∼880 Å^2^, respectively) structures [37], providing further evidence of the structural independence of MosV-RBP and NarV-RBP from structurally characterized paramyxoviral RBPs.

Furthermore, we note that the observed mode of MosV-RBP_β_ and NarV-RBP_β_ dimerization is in agreement with the location of N-linked glycosylation sequons (NXS/T where X≠P). Indeed, although MosV-RBP_β_ and NarV-RBP_β_ lack the high level of glycosylation inherent to most paramyxoviral proteins [42, 44-50], in line with the hypothesis that glycosylation is not expected to be occluded within protein–protein interfaces, the single N-linked sites observed in the β-propeller domain of both MosV-RBP (Asn319) and NarV-RBP (Asn575) are directed away from and do not interfere with the observed RBP homodimers. Furthermore, we note that the MosV/NarV exhibit the greatest level of structural conservation in the region of this oligomeric interface with respect to the rest of the molecule (**Fig. 2B**), suggestive that these blades of the β-propeller are structurally constrained to promote a similar mode of overall assembly.

To determine the oligomeric state of glycosylated NarV-RBP_β_ and MosV-RBP_β_ in solution, we performed size-exclusion chromatography-coupled small-angle X-ray scattering (SEC-SAXS) on the glycosylated proteins (**Table S3, Fig. S2-S4**). Under-dilute conditions, SAXS measures the shape and size of macromolecular particles in solution [51]. SAXS measurements suggest both proteins are compact, globular proteins in solution (Figure S4A). For each protein, fitting of the monomeric RBP_β_ structures to either SEC-SAXS datasets was exceptionally poor (χ^2^=80.0 and χ^2^=105.6 for NarV-RBP_β_ and MosV-RBP_β._ respectively) suggesting that their monomeric forms do not represent the solution state. In addition, model independent analysis of the respective pair-distance distribution functions (Figure S4B) for NarV-RBP_β_ and MosV-RBP_β_ shows a shoulder at ∼1/2 maximum height at 60 to 70 Å suggesting a dimeric form of the protein. Significant improvements in the fits to the SAXS curves were observed when using the crystallographic dimeric forms (χ^2^=1.95 and χ^2^=4.50 for NarV-RBP_β_ and MosV-RBP_β._ respectively). However, the crystallographic models are incomplete, as electron density for the N- and C-terminal residues and GlcNAc_2_Man_9_ glycans (derived by kifunensine treatment) were not observed. To complete the models, we used simulated annealing molecular dynamics (SA-MD) simulations with torsion angle restraints and additional hydrogen-bond and distance restraints derived from the respective crystal structures. The high temperature simulated annealing was cycled repeatedly producing ∼1,000 sampled conformations of each RBP_β_ protein. Using this SA-MD approach, a set of best fitting models was identified, which demonstrated an expected flexibility across the glycans and disordered termini. Using the updated dimeric forms of the structures, the fits to the SAXS curves were improved to χ^2^= 0.65 and χ^2^=1.28, indicative that the derived models represent the solution state of NarV-RBP_β_ and MosV-RBP_β._ respectively. The resulting MosV-RBP_β_ and NarV-RBP_β_ dimers retained angular differences in subunit association that were in line with the values of the dimeric crystal structures, 81° and 89°, respectively (**Fig. S4**). Additionally, the MosV-RBP_β_ and NarV-RBP_β_ best fit structures generated by high temperature simulated annealing, largely retained the extensive homodimeric interfaces found in both crystal structures. Post-MD, MosV-RBP_β_ and NarV-RBP_β_ retained ∼1,370 Å^2^ and ∼1,880–1,960 Å^2^ (calculated from 3 of the best fitting NarV-RBP MD models) of the previously calculated ∼2,070 Å^2^ and ∼2,170 Å^2^ buried surface area, respectively. These results are summarized in **Supplemental Table S3**. This analysis indicates that the RBP_β_ dimerization occurs in solution, further supporting the hypothesis that our structurally observed homodimeric narmovirus RBP_β_ organization resembles a biologically relevant assembly (**Fig. 4**). Furthermore, this analysis also supports the intrinsic flexibility of surface-exposed loop regions that exhibit high RMSD values upon overlay of NarV-RBP_β_ and MosV-RBP_β_ crystal structures (**Fig. 2B**). *In toto*, this integrated structure, solution state analysis, and molecular dynamics approach indicates that our structurally-observed homodimeric interfaces of narmovirus RBPs are unlikely to be features that are solely specific to crystallization.

**Figure 4.**
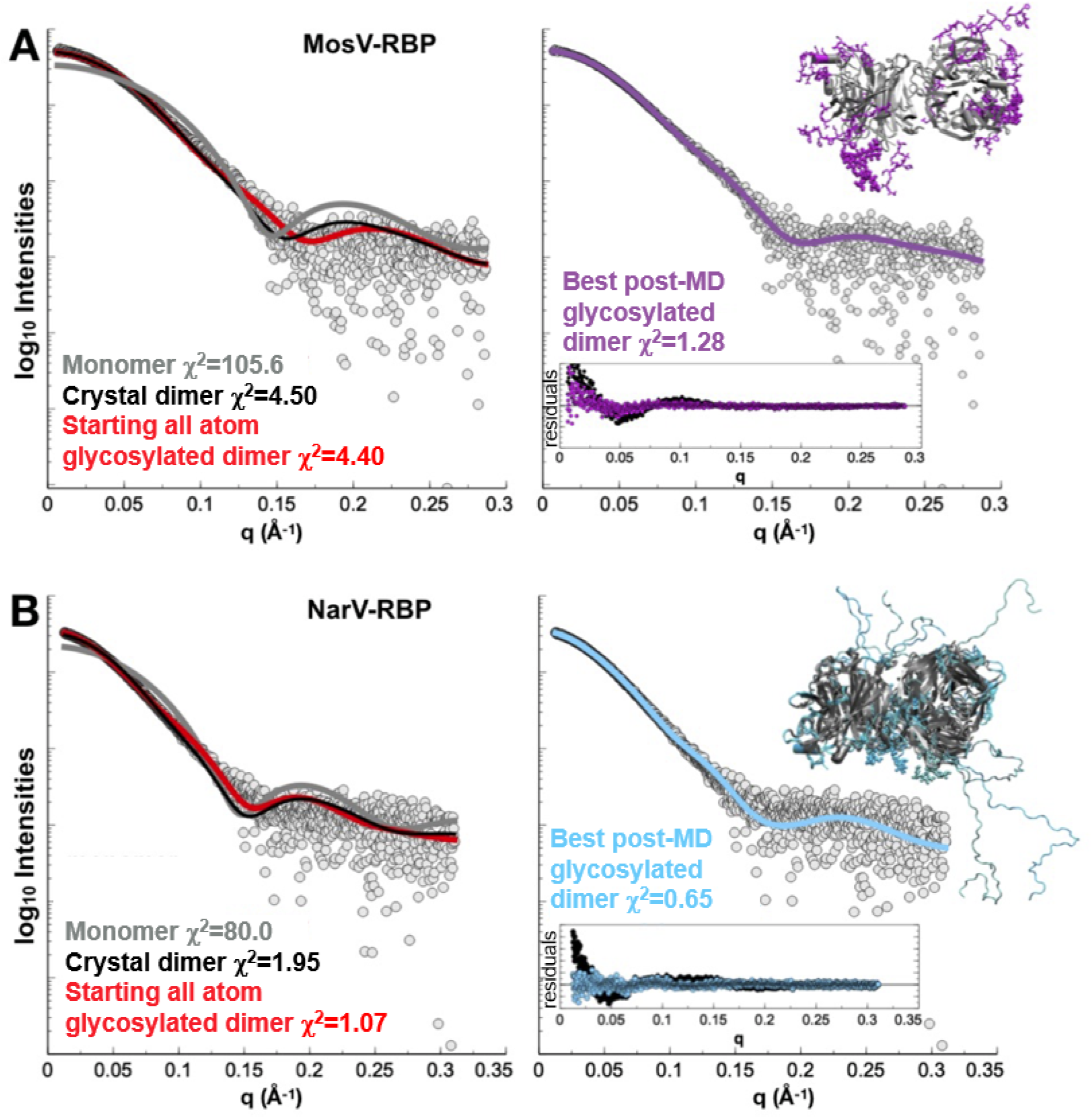
SAXS analysis validates the organization of MosV-RBP_β_ and NarV-RBP_β_ as dimers in solution. Experimental MosV-RBP (A) and NarV-RBP (B) SAXS data (grey circles) compared to theoretical SAXS curves calculated from crystallographically observed monomeric (dark grey line) and, dimeric (black line) oligomeric states. In both cases, crystallographic structure determination did not reveal electron densities for N-, C-termini and glycosylations that were present during the SAXS experiment. The SAXS datasets fit better to completed structures (addition of modelled N- and C-termini, and glycosylations, red) than the crystal structures. Application of high-temperature, simulated annealing molecular dynamics (MD) of completed, dimeric structures (purple and cyan for, MosV-RBP_β_ and NarV-RBP_β_, respectively) improved fitting further. Log intensity (I(q)) is plotted against the Porod invariant (q) [66]. χ^2^ values are shown and were generated by calculating the fit between structural models (*i*.*e*., ‘Monomer’, ‘Crystal dimer’, ‘Starting all atom glycosylated dimer’, and ‘Best post-MD glycosylated dimer’) and the experimental (SAXS) data. The best χ^2^ values were obtained following MD of glycosylated, all atom MosV-RBP (χ^2^=1.28) and NarV-RBP (χ^2^=0.65) dimers. Inset plots (right) show the residuals calculated against the final models.

### Narmoviral RBPs lack motifs associated with known modes of paramyxovirus entry

Structural studies of H, HN, and G-type RBPs have revealed conserved features that confer shared receptor specificity [3, 4, 7, 27, 31-35, 52, 53]. For example, with the exception of MojV, which undergoes a host-cell entry pathway distinct from other characterized henipaviruses [29], henipaviral G RBPs exhibit increased levels of sequence conservation at known ephrin binding sites [3, 4, 27, 31, 53]. Similarly, morbillivirus H RBPs exhibit elevated levels of sequence conservation at SLAMF1 and nectin-4 receptor-binding sites [7, 52], and HN glycoproteins encode well-conserved motifs associated with hemadsorption and neuraminidase activity [32-35].

In agreement with the structural distinctiveness of narmovirus RBPs (**Fig. 2**), MosV-RBP_β_ and NarV-RBP_β_ lack features associated with receptor recognition by G, H, and HN-type RBPs (**Fig. 5**). Firstly, mapping analysis reveals a low level of sequence conservation between our narmovirus RBP structures and previously reported henipaviral RBPs at characterized ephrin receptor binding sites (∼26%–28% sequence identity) (**Fig. 5A**). Furthermore, overlay of henipaviral RBP-ephrinB1 (6THG) [27, 31], -ephrinB2 (2VSM) [3], and -ephrinB3 (3D12) [4] structures onto our MosV-RBP_β_ and NarV-RBP_β_ revealed substantial clashes and indicate that MosV-RBP_β_ and NarV-RBP_β_ lack the hydrophobic pockets required to accommodate the G–H loop of B-type ephrin ligands [3, 4, 27, 31]. In particular, we note that recognition would likely be sterically precluded by the extended β5L01 loop of MosV-RBP_β_ and NarV-RBP_β_, which is not present in the HNV-G RBPs (**Supplementary Fig. S5**) [3, 4, 27, 31, 53]. Secondly, MosV-RBP_β_ and NarV-RBP_β_ maintain low levels of sequence identity with MeV-H RBP [26] at SLAMF1 (12% and 9% sequence identity, respectively) and nectin-4 (7% and 10% sequence identity, respectively) binding sites (**Fig. 5B**), and further, lack the β4-5 groove integral to morbilliviral receptor interactions [25, 26]. Thirdly, the narmoviral RBPs lack many of the residues that are central to the sialic acid interacting functionality of HN-RBPs (**Fig. 5C**), including only 1–4 of the seven conserved sialidase residues (Arg_1_, Asp_1_, Glu_4_, Arg_4_, Arg_5_, Tyr_6_, and Glu_6_), and only 1–2 residues of the hexapeptid motif (‘Asn–Arg–Lys–Ser–Cys–Ser’) [54]. Additionally, despite being present in the primary amino acid sequence of both MosV-RBP and NarV-RBP, structural analysis reveals that Tyr_6_ is not presented within the putative sialic acid binding site. The presence of a high level of variability in this region of the molecule is suggestive that narmoviruses lack the conserved receptor-specific functionality found in HN RBPs [32-36]. Combined, this comparative analysis demonstrates that MosV-RBP and NarV-RBP lack the features known to be associated with characterized H, HN, and G-type receptor-mediated host-cell recognition pathways.

**Figure 5.**
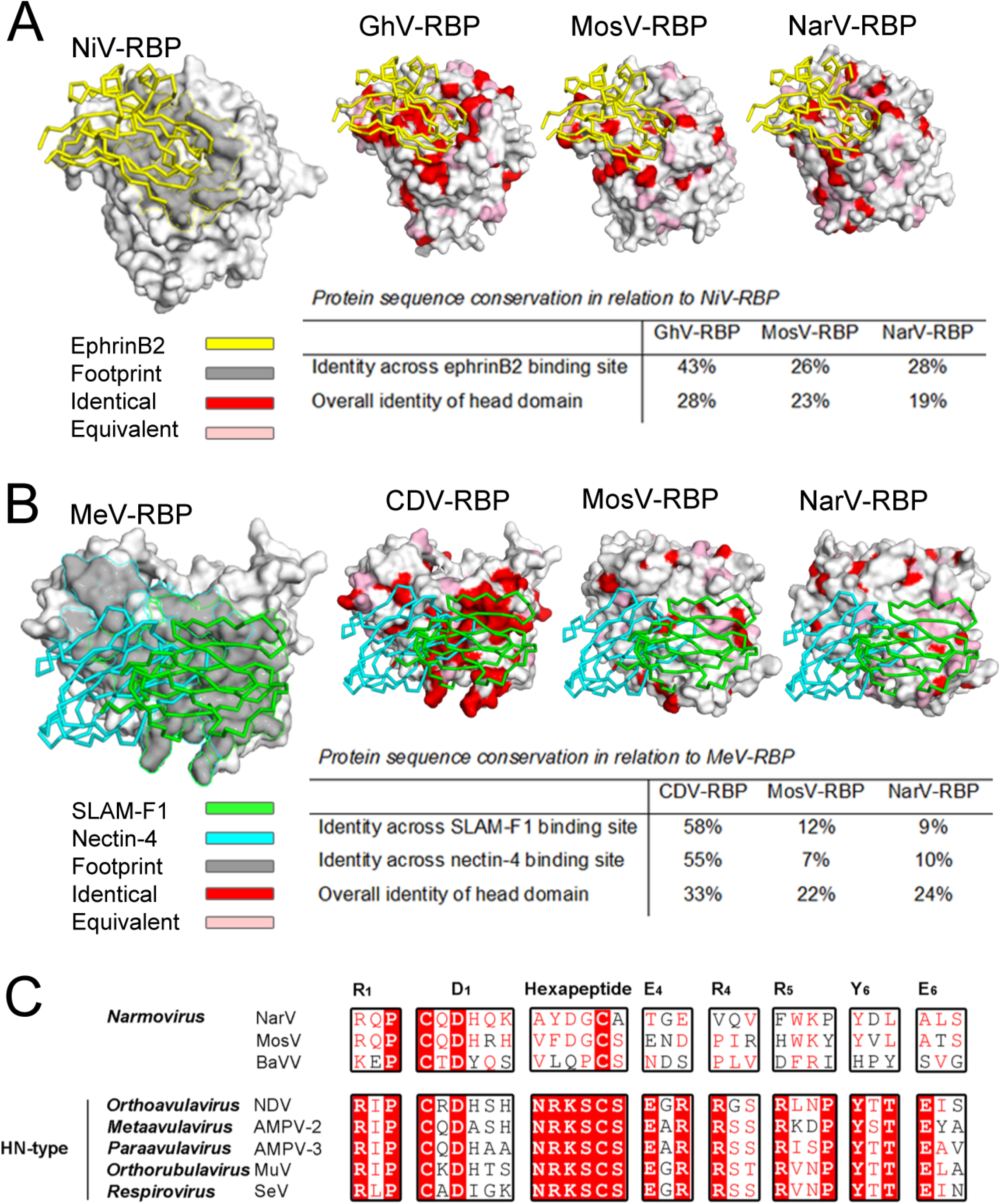
MosV-RBP and NarV-RBP lack motifs associated with known modes of paramyxovirus entry. (**A**) NiV-RBP (PDB ID 2VSM) [3] is shown as a surface with ephrinB2 shown as a yellow ribbon and the cognate receptor-binding footprint coloured grey and outlined in yellow (far left). Ghana virus (GhV) RBP (second left; 4UF7), MosV-RBP (second right), NarV-RBP (far right) are shown in surface representation with residues coloured according to sequence conservation with NiV-RBP. Identical residues are colored red and equivalent pink. (**B**) MeV-RBP (PDB ID 3ALZ) [26] is shown in surface representation with SLAM-F1 (green) and nectin-4 (cyan) shown in ribbon representation. The cognate joint receptor-binding footprint is colored grey (far left) and outlined according to the SLAM-F1 (green) and nectin-4 (cyan) binding footprints. As no CDV-RBP structure is currently available, sequence conservation of CDV-RBP (second left) with MeV-RBP is mapped onto the surface of MeV-RBP. MosV-RBP (second right) and NarV-RBP (far right) are shown in surface representation with residues colored as above, according to sequence conservation with MeV-RBP. (**C**) Alignment of the RBP amino acid sequences from MosV (NP_958054.1), NarV (YP_006347588.1), BaVV (ATW63189.1), Newcastle disease virus (NDV) (YP_009512963.1), AMPV-2 (AQQ11616.1), AMPV-3 (AWU68199.1), Mumps virus (MuV) (MG460606.1) and Sendai virus (SeV) (NC_001552.1). The seven conserved sialidase residues [36] and hexapeptide motif [54] are labelled according to residue and blade location [36] and annotated above alignments.

## Discussion

Viral genome sequencing within wildlife reservoirs has provided glimpses into the extensive genetic diversity that exists within the *Paramyxoviridae*, allowing expansion of the number of genera within the family [2]. Rodent-borne paramyxoviruses from the genus *Narmovirus* exemplify this diversity, where the founding members, NarV and MosV (from Trinidad and Tobago and Australia, respectively), have been putatively joined by recently identified narmoviruses from the UK [13] and Russia [12]. Indeed, the growth of this genus reflects our increasing appreciation of the broad geographic range assumed by this group of pathogens. However, despite this surprising prevalence, little is known about narmovirus pathobiology, host-range, or inter-species transmission potential, features that are expected to be modulated, at least in part, by the RBP [1, 7].

Here, we take an initial step to address this paucity of knowledge through structural determination of the C-terminal head region of the RBP from NarV and MosV. Our investigation reveals that whilst the head domains of these narmovirus RBPs retain the six-bladed β-propeller fold observed across paramyxovirus RBPs, they lack the residues and complimentary surfaces required for known paramyxoviral receptor interactions. Indeed, both NarV-RBP_β_ and MosV-RBP_β_ do not present the characteristic ephrinB-class binding pocket presented by the henipaviral RBPs [3, 4] or the β4–β5 groove integral to the interaction between MeV and its receptors [24-26]. Furthermore, they also lack many of the stringently conserved residues associated with sialic acid binding and hydrolyzing activity [32-36].

In previous structural studies, we and others have shown that structure-based phylogenetic analysis can classify viral proteins according to conformational state and receptor specificity [22, 29, 37, 41, 55, 56]. Consistent with the observation that narmovirus RBPs lack known paramyxovirus receptor-specific features (**Fig. 3**), our structure-based phylogenetic analysis reveals that NarV and MosV form a distinct structural class that is nearly equidistant from known H, HN, and G RBP structures (**Fig. 4**). Furthermore, by analogy to the large structural difference between ephrin-binding HNV-G RBPs and the functionally distinct MojV-G RBP [29], NarV and MosV RBP exhibit large RMSDs upon structural overlay with reported paramyxoviral RBP structures (≥2.1–3.3 Å).

Finally, NarV-RBP_ß_ and MosV-RBP_ß_ were observed to be dimeric in solution (**Fig. 4**). This crystallographically observed and SAXS validated angle of dimeric organization for NarV-RBP_ß_ and MosV-RBP_ß_ (∼90°) contrasts that observed for sialic acid-binding HN-type RBPs (∼60°) [16, 32-35]. Divergence from this angle has also been observed in protein-specific HeV-G (∼40°) and MeV-H (∼120°) RBD structures, and has been hypothesized to have arisen out of a requirement to recognize different, potentially larger receptors [3, 23-26, 57]. These combined observations support a model whereby NarV and MosV undergo a host-cell entry pathway that is distinct from characterized H, HN, and G RBP-bearing paramyxoviruses.

In sum, our NarV-RBP_ß_ and MosV-RBP_ß_ structures provide molecular-level blueprints that support the independent functional and structural classification of narmovirus RBPs from other paramyxoviruses [2]. This work offers a platform for future investigations focused on assessing and rationalizing the pathobiological characteristics and the receptor(s) utilized by narmoviruses for host-cell entry. While the threat that NarV, MosV, and other narmoviruses pose to human health and animal husbandry remains unknown, by defining the RBP architecture assumed by this group of viruses, this work renders us better prepared to understand and respond to the emergence of pathogenic narmoviruses, if they emerge.

## Materials and Methods

### Protein production

MosV and NarV RBP constructs for protein expression were generated from human codon-optimized genes of the full-length RBPs (Genbank AY286409.1 and FJ362497.2, respectively). For crystallization, MosV-RBP_β_ (Glu202–Thr632) and NarV-RBP_β_ (Glu184–Pro657) were cloned into the pHLsec mammalian expression vector, which encodes a C-terminal hexahistidine tag [58]. For functional studies, MosV-RBP (Thr157–Thr632) and NarV-RBP (Glu184–Pro657) were cloned into the pHLsec mammalian expression vector encoding a C-terminal Fc tag [58]. No soluble protein was obtained following the addition of a C-terminal Fc-tag to the MosV-RBP_β_ (Glu202–Thr632) construct, therefore it was deemed necessary to extend the N-terminal construct boundary for MosV-RBP to produce this tagged protein.

Protein was produced by transient transfection of human embryonic kidney (HEK) 293T cells, and secreted protein was harvested after 72 h incubation at 37 °C, 5% CO_2_. For crystallization and SAXS analysis, protein was produced in the presence of 5 µM kifunensine [38] and purified using immobilized metal-affinity chromatography (IMAC). Prior to application on the HisTrap™ HP (Cytvia) column, cell supernatant was exchanged into 10 mM Tris (pH 8.0), 150 mM NaCl and concentrated using an ÄKTA Flux diafiltration system (Cytvia). His-tagged protein was eluted using 250 mM imidazole. Subsequently, if required for crystallization, N-linked sugars were cleaved at the di-N-acetylchitebiose core using endoglycosidase F1 (EndoF1) (10 µg/mg protein, 12 h, 21 °C). Size-exclusion chromatography (SEC) was performed in 10 mM Tris pH 8.0, 150 mM NaCl buffer using a Superdex™ 200 10/30 column (Cytvia). For cell-binding analysis, cell supernatants were exchanged into 20 mM sodium phosphate pH 7.0 buffer prior to affinity purification on a HiTrap® Protein G HP (Cytvia) column. Fc-tagged protein was eluted by washing the column into 0.1 M glycine-HCl pH 2.7 before immediate neutralization with 60 µL of 1 M Tris-HCl pH 9.0 per mL of eluate. Subsequently, Fc-tagged protein was purified and exchanged into 20 mM sodium phosphate pH 7.0 buffer using SEC on a Superdex™ 200 10/30 column (Cytvia).

### Cell-binding studies

Vero cells were maintained in DMEM with 10% heat inactivated (HI) Fetal Bovine Serum (FBS). CHO pgsA745 hamster cells were maintained in DMEM/F12 medium supplemented with 10% HI FBS. LA-4 cells, a mouse lung epithelial cell lines, were obtained from the ‘American Type Culture Collection’ (ATCC) and maintained in Ham’s F12K medium supplemented with 15% HI FBS. Cultured cells were collected with 10 mM EDTA, then incubated for 1hr with soluble, Fc-tagged receptor binding protein. Cells were then washed twice in 2% FBS in Dulbecco’s phosphate buffered saline (DPBS), stained with secondary anti-Fc-APC (allophycocyanin) antibody at a 1:2,000 dilution and washed twice again. Cells were subsequently fixed in 2% paraformaldehyde and resuspended in 2% FBS in DPBS prior to flow cytometry (Guava easyCyte).

### Crystallization and structure determination

MosV-RBP_β_ crystals were grown using the nanoliter-scale sitting-drop vapor-diffusion method at room temperature, using 100 nl protein (5.5 mg/mL) and 100 nl reservoir [59]. Crystals grew in a precipitant containing 0.2 M L-arginine, 0.1 M Tris pH 7.8, 8% poly-γ-glutamic acid (PGA)-LM, 6% dextran sulphate and were immersed in 20% glycerol prior to cryo-cooling by plunging into liquid nitrogen. Initial X-ray diffraction data were collected at a wavelength of 0.9795 Å on beamline I03, Diamond Light Source (DLS). Reflections were processed to a resolution of 2.75 Å using the xia2 package [60] (**Table S1**). For experimental phasing, crystals from this same condition were soaked in a solution containing potassium tetrachloroplatinate (K_2_PtCl_4_) for 3 h, prior to cryo-cooling with a 20% glycerol solution containing K_2_PtCl_4_ diluted in precipitant. Crystals were exposed to a wavelength consistent with the platinum LIII absorption edge (λ = 1.072 Å) at beamline I04, DLS. The isomorphous differences between the native dataset and the derivatized dataset, in addition to the anomalous signal derived from platinum, enabled subsequent phase determination using single isomorphous replacement with anamolous scattering (SIRAS) using the Autosol wizard [40]. A further MosV-RBP_β_ dataset was collected on crystals which grew in the following precipitant mixture: 0.2 M potassium bromide, 0.2 M potassium thiocynate, 0.1 M sodium cacodylate pH 6.5, 3% PGA-LM, 20% w/v polyethylene glycol (PEG) 550 MME. Crystals were harvested and cryo-cooled in a 20% glycerol solution diluted with precipitant. Data were collected at a wavelength of 0.9795 Å, on beamline I04, DLS, and reflections were processed to 1.62 Å using the xia2 package [60] (**Table 1**). Molecular replacement using PHASER [61] with the initial MosV-RBP model was utilized to solve the higher-resolution dataset.

NarV-RBP_β_ crystals were grown using nanolitre-scale sitting-drop vapor-diffusion at room temperature, using 100 nl protein (4.5 mg/mL) and 100 nl reservoir [59]. Small crystals grew in a precipitant containing 0.2 M magnesium chloride, 0.1 M HEPES pH 7.5, 25% PEG 3350, 10% PEG 400. Crystals were optimized by seeding into a lower concentration solution of NarV-RBP_β_ (Bergfors 2007; Walter et al. 2008). Crystals were pulverized, utilizing a seed bead (Hampton Research, USA) (Luft and DeTitta 1999), and dispensed onto a plate prepared with the original precipitant mix and NarV-RBP_β_ at a concentration of 3 mg/mL (Walter et al. 2008). A crystal was immersed in 20% glycerol prior to cryo-cooling by immersion into liquid nitrogen. Data were collected at a wavelength of 0.9795 Å on beamline I04, DLS, and reflections were processed to 2.07 Å using the xia2 package [60] (**Table S1**). A partially refined model of MosV-RBP was used to solve the structure of NarV-RBP_β_ by molecular replacement with PHASER [61]. For all models, building and structure refinement were iteratively performed using the programmes COOT and Phenix.Refine, respectively (**Table S2**) [62, 63]. Non-crystallographic symmetry (NCS) restraints were employed throughout, and translation-libration-screw (TLS) parameters employed for later rounds of refinement. Models were validated using the Molprobity server [64].

### SAXS with in-line high performance liquid chromatography

SAXS was used to characterize the solution state of glycosylated MosV-RBP_β_ and NarV-RBP_β_ (**Table S3**). Data were collected on beamline B21, at the DLS, configured to measure across the scattering vector range 0.0032 Å-1 < q < 0.38 Å-1. For high performance liquid chromatography (HPLC) mode, a 45 µL sample at a concentration of 5 mg/mL was loaded onto a Superdex™ 200 PC 3.2/30 (Cytvia). Buffer TBS, was washed over at a rate of 0.075 mL/min. The SAXS instrument was coupled directly with inline SEC with exposures collected every 2 seconds [65]. SEC-SAXS profiles corresponding to a single chromatographic separation were analyzed with the program, ScÅtter, where peak and background selection and data reduction (www.bioisis.net) were performed to produce a single SAXS curve for each protein sample.

### Molecular dynamics with SAXS

Molecular dynamics simulations were performed with the crystallography and NMR systems (CNS) program CNSsolve version 1.3 (http://cns-online.org/v1.3/). Missing N- and C-terminal tails were added back using the generate_seq.inp, generate.inp and model_anneal.inp script from CNS. Crystallographic structures for each RBP served as templates to derive NOE like distance restraints that folded a starting extended polypeptide chain into the respective, folded, crystallographic monomeric RBP. Scale factors for the NOE energy term and molecular dynamics time steps were adjusted down to minimize large energy terms in the gradient descent. For each refold, greater than 20 models were produced from independent random starts. The model with the lowest energy term (maximized NOE-like distance restraints) served as the base template for further model building. Base templates were then uploaded to the GLYCAM-Web server (https://glycam.org) to add the GlcNAc_2_Man_9_ glycans. The glycosylated monomer was duplicated and superimposed onto the crystallographic subunits to produce a full-length, glycosylated RBP dimer for each virus. The completed models were then used with model_anneal.inp for simulated annealing (SA), torsion angle MD to sample conformation space. Sampling was performed using randomly selected distance restraints (∼3% non-hydrogen, backbone and C-beta distance pairs) derived from the crystallographic models. Residues with B-factors greater than 2-times the average were excluded from selection, which coincided with residues in the loops forming the rim of the β-propeller. Additional intersubunit hydrogen bond restraints were derived between the subunit interfaces to act as rigid restraints. The upper and lower limits for each restraint was set by the Shannon resolution (л/q_max_). Several rounds of SA-MD produced a set of refolded models that varied slightly in the conformation of the sugars, loops and N- and C-termini. SAXS profiles were calculated for MosV-RBP_β_ and NarV-RBP_β_ structures using the fast X-ray scattering (FoXS) server [66], and matched to SAXS experimental profiles. Determination of the success of the fit was demonstrated by assessment of the χ^2^ statistic as well as the FOXS parameters c_1_ and c_2,_ with c_2_ < 4 (c_2_ > 4 suggests over-fitting). **Structure-based phylogenetic analysis**. For structural phylogenetic analysis, RBP six-bladed β-propeller monomers were prepared by removal of water molecules, ligands, and protein residues outside of the canonical fold. Structures were analyzed with the Structural Homology Program, SHP [67, 68]. Pairwise evolutionary distance matrices were used to generate un-rooted phylogenetic trees in PHYLIP [69].

### Dimer angle analysis

UCSF Chimera was utilized to analyze relative angles of monomers within the paramyxoviral dimers [70]. Planes representing the top faces of the monomer subunits were constructed based upon conserved stretches of paramyxoviral RBP sequence using the ‘Define plane functionality’, with the angle between the monomers of a dimer calculated using these planes.

### Data deposition

The atomic coordinates and structure factors for MosV-RBP_β_ and NarV-RBP_β_ have been deposited in the Protein Data Bank with the accession codes 7ZM5 and 7ZM6, respectively. MosV-RBP_β_ and NarV-RBP_β_ SAXS datasets have been deposited in bioISIS.net with accession codes MSVRB1 and NRVRB1.

## Acknowledgements

We are grateful to DLS for beamtime (proposals mx10627 and mx14744) and the staff of beamlines I02, I04, and B21 for assistance with data collection. We thank the Medical Research Council (MR/L009528/1 and MR/S007555/1 to T.A.B.), Engineering and Physics Research Council (EP/K503113/1, EP/L505031/1, EP/M50659X/1 and EP/M508111/1 to A.S), Academy of Finland (#309605 to I.R.), and NIH (NIAID AI123449, AI069317, and AI115226 to B.L.) for funding. B.L. also acknowledges the Ward Coleman estate for endowing the Ward-Coleman Chairs at the Icahn School of Medicine at Mount Sinai. The Wellcome Centre for Human Genetics is supported by Wellcome Centre grant 203141/Z/16Z.

